# Biodiversity Soup II: A bulk-sample metabarcoding pipeline emphasizing error reduction

**DOI:** 10.1101/2020.07.07.187666

**Authors:** Chunyan Yang, Kristine Bohmann, Xiaoyang Wang, Wang Cai, Nathan Wales, Zhaoli Ding, Shyam Gopalakrishnan, Douglas W. Yu

## Abstract

1. Despite widespread recognition of its great promise to aid decision-making in environmental management, the applied use of metabarcoding requires improvements to reduce the multiple errors that arise during PCR amplification, sequencing, and library generation. We present a co-designed wet-lab and bioinformatic workflow for metabarcoding bulk samples that removes both false-positive (tag jumps, chimeras, erroneous sequences) and false-negative (‘dropout’) errors. However, we find that it is not possible to recover relative-abundance information from amplicon data, due to persistent species-specific biases.
2. To present and validate our workflow, we created eight mock arthropod soups, all containing the same 248 arthropod morphospecies but differing in absolute and relative DNA concentrations, and we ran them under five different PCR conditions. Our pipeline includes qPCR-optimized PCR annealing temperature and cycle number, twin-tagging, multiple independent PCR replicates per sample, and negative and positive controls. In the bioinformatic portion, we introduce *Begum*, which is a new version of *DAMe* (Zepeda-Mendoza *et al*. 2016. *BMC Res. Notes* 9:255) that ignores heterogeneity spacers, allows primer mismatches when demultiplexing samples, and is more efficient. Like *DAMe, Begum* removes tag-jumped reads and removes sequence errors by keeping only sequences that appear in more than one PCR above a minimum copy number per PCR. The filtering thresholds are user-configurable.
3. We report that OTU dropout frequency and taxonomic amplification bias are both reduced by using a PCR annealing temperature and cycle number on the low ends of the ranges currently used for the Leray-FolDegenRev primers. We also report that tag jumps and erroneous sequences can be nearly eliminated with *Begum* filtering, at the cost of only a small rise in dropouts. We replicate published findings that uneven size distribution of input biomasses leads to greater dropout frequency and that OTU size is a poor predictor of species input biomass. Finally, we find no evidence for ‘tag-biased’ PCR amplification.
4. To aid learning, reproducibility, and the design and testing of alternative metabarcoding pipelines, we provide our Illumina and input-species sequence datasets, scripts, a spreadsheet for designing primer tags, and a tutorial.

## Introduction

DNA metabarcoding enables rapid and cost-effective identification of taxa within biological samples, combining amplicon sequencing with DNA taxonomy to identify multiple taxa in bulk samples of whole organisms and in environmental samples such as water, soil, and feces (Taberlet *et al*. 2012a; Taberlet *et al*. 2012b; Deiner *et al*. 2017). Following initial proof-of-concept studies (Fonseca *et al*. 2010; Hajibabaei *et al*. 2011; Thomsen *et al*. 2012; Yoccoz 2012; Yu *et al*. 2012; Ji *et al*. 2013) has come a flood of basic and applied research and even new journals and commercial service providers (Murray, Coghlan & Bunce 2015; Callahan *et al*. 2016; Zepeda-Mendoza *et al*. 2016; Alberdi *et al*. 2018; Zizka *et al*. 2019). Two recent and magnificent surveys are Taberlet *et al*. (2018) and Piper *et al*. (2019). The big advantage of metabarcoding as a biodiversity survey method is that with appropriate controls and filtering, metabarcoding can estimate species compositions and richnesses from samples in which taxa are not well characterized *a priori* or reference databases are incomplete or lacking. However, this is also a disadvantage, because we must first spend effort to design reliable and efficient metabarcoding pipelines.

Practitioners are thus confronted by multiple protocols that have been proposed to avoid and mitigate the many sources of error that can arise in metabarcoding (Table 1). These errors can result in false negatives (failures to detect target taxa that are in the sample, ‘*dropouts*’), false positives (false detections of taxa), poor quantification of biomasses, and/or incorrect assignment of taxonomies, which also results in paired false negatives and positives. As a result, despite recognition of its high promise for environmental management (Ji *et al*. 2013; Hering *et al*. 2018; Abrams *et al*. 2019; Bush *et al*. 2019; Piper *et al*. 2019; Cordier *et al*. 2020; Cordier 2020), the applied use of metabarcoding is still getting started. A comprehensive understanding of costs, the factors that govern the efficiency of target taxon recovery, the degree to which quantitative information can be extracted, and the efficacy of methods to minimize error is needed to optimize metabarcoding pipelines (Hering *et al*. 2018; Axtner *et al*. 2019; Piper *et al*. 2019).

**Table 1.**
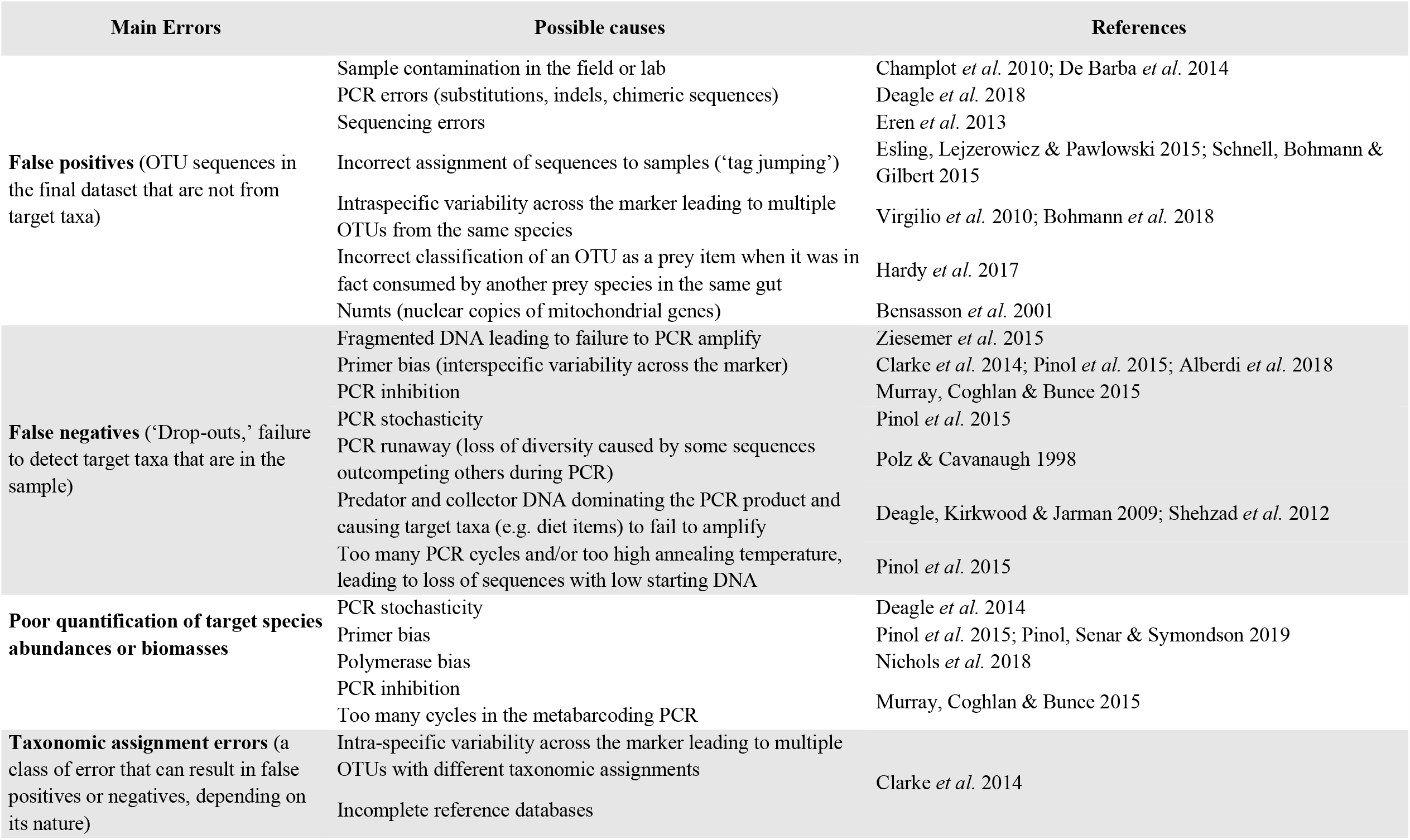
Four classes of metabarcoding errors and their causes. Not included are software bugs, general laboratory and field errors like mislabeling, sampling biases, or inadequate sequencing depth.

Here we consider one of the two main sample types used in metabarcoding: bulk-sample DNA (the other type being environmental DNA, Bohmann et al., 2014). Bulk-sample metabarcoding, such as mass-collected invertebrates, is being studied as a way to generate multi-taxon indicators of environmental quality (Lanzén *et al*. 2016; Hering *et al*. 2018), to track ecological restoration (Cole *et al*. 2016; Fernandes *et al*. 2018; Barsoum *et al*. 2019; Wang *et al*. 2019), to detect pest species (Piper *et al*. 2019), and to understand the drivers of species diversity gradients (Zhang *et al*. 2016).

We present a co-designed wet-lab and bioinformatic pipeline that uses qPCR-optimized PCR conditions, three independent PCR replicates per sample, twin-tagging, and negative and positive controls to: (i) remove sequence-to-sample misassignment due to tag-jumping, (ii) reduce dropout frequency and taxonomic bias in amplification, and (iii) reduce false-positive frequency.

As part of the pipeline, we introduce a new version of the *DAMe* software package (Zepeda-Mendoza *et al*. 2016), renamed *Begum* (Hindi for ‘lady’), to demutiplex samples, remove tag-jumped sequences, and filter out erroneous sequences (Alberdi *et al*. 2018). Regarding the latter, the *DAMe*/*Begum* logic is that true sequences are more likely to appear in multiple, independent PCR replicates and in multiple copies than are erroneous sequences (indels, substitutions, chimeras). Thus, erroneous sequences can be filtered out by keeping only sequences that appear in more than one (or a low number of) PCR replicate(s) at above some minimum copy number per PCR, albeit at a cost of also filtering out some true sequences. *Begum* improves on *DAMe* by ignoring heterogeneity spacers in the amplicon, allowing primer mismatches during demultiplexing, and by being more efficient. We note that this logic is less applicable to species represented by trace DNA, such as in filtered water samples, where low concentrations of DNA template are more likely to result in a species truly appearing in only one PCR (Piaggio *et al*. 2014; Harper *et al*. 2018).

To test our pipeline, we created eight ‘mock’ arthropod soups, each consisting of the DNA of the same 248 arthropod taxa mixed together in the lab and differing in absolute and relative DNA concentrations, ran them under five different PCR conditions, and used *Begum* to filter out erroneous sequences (Fig.1). We then quantified the efficiency of species recovery from bulk arthropod samples, as measured by four metrics:

1. the frequency of false-negative OTUs (‘dropouts’, i.e. unrecovered input species),
2. the frequency of false-positive OTUs (sequences not from the input species),
3. the recovery of frequency information (does OTU size [number of reads] predict input DNA amount per species?), and
4. taxonomic bias (are some taxa more or less likely to be recovered?).

Highest efficiency is achieved by recovering *all* and *only* the input species, in their original frequencies. We show that with *Begum* filtering, metabarcoding efficiency is highest with a PCR cycle number and annealing temperature at the low ends of the ranges currently used in metabarcoding studies, that *Begum* filtering nearly eliminates false-positive OTUs, at the cost of only a small absolute rise in dropout frequency, that greater species evenness and higher concentrations reduce dropouts (replicating Elbrecht, Peinert & Leese 2017), and that OTU sizes are not reliable estimators of species-biomass frequencies. We also find at most only small effect sizes for ‘tag bias,’ which is the hypothesis that the sample-identifying nucleotide sequences attached to PCR primers might promote annealing to some template-DNA sequences over others (e.g. Berry *et al*. 2011; O’Donnell *et al*. 2016), exacerbating taxonomic bias in PCR. All these results have important implications for using metabarcoding as a biomonitoring tool.

## Methods

In S06_Extended Methods, we present an unabridged version of this Methods section.

### Mock soup preparation

#### Input species

– We used Malaise traps to collect arthropods in Gaoligong Mountain, Yunnan province, China. From these, we selected 282 individuals that represented different morphospecies, and from each individual, we separately barcoded DNA from the leg and the body. After clustering, we ended up with 248 97%-similarity DNA barcodes, which we used as the ‘input species’ for the mock soups (S07_MTBFAS.fasta).

#### COI and genomic DNA quantification

– To create the eight mock soups with different concentrations evenness of the 248 input species, we quantified DNA concentrations of their legs and bodies, using qPCR and a reference standard-curve method on the QuantStudio 12K Flex Real-Time PCR System (Life Technologies, Singapore) with Leray-FolDegenRev primers (Yu *et al*. 2012; Leray *et al*. 2013). We then diluted each species to their target DNA concentrations (Tables 2, S03). After dilution, we also measured each species’ genomic DNA concentrations, to test whether OTU size can predict genomic DNA, which is a proxy measure for animal biomass.

**Table 2.** The eight mock soups, each containing the same 248 arthropod OTUs but differing in absolute (Body/Leg) and relative (*Hhml, hhhl, hlll*, and *mmmm*) DNA concentrations. Numbers in the table are the numbers of OTUs in each concentration category (*H, h, m, l*). Thus, the *Hhml_body* soup contains 50 species with a DNA concentration between 50-200 ng/μl, each added as an aliquot of 1 μl, and so on. The evenness of DNA concentrations in each mock soup is summarized by the Shannon index. Higher values indicate a more even distribution. A few species provided only a low level of DNA concentration but were included in the *mmmm* soup as such.

| DNA extraction<br>from arthropod<br>body part | DNA<br>concentration<br>evenness | Number of OTUs in each concentration category |  |  |  | Total number of<br>OTUs | Shannon index |
| --- | --- | --- | --- | --- | --- | --- | --- |
|  |  | High (*H*) | high (*h*) | medium (*m*) | low (*l*) |  |  |
|  |  | 50-200 ng/μl | 10-48 ng/μl | 1-8 ng/μl | 0.001-0.1 ng/μl |  |  |
| Body | *Hhml* | 50 | 75 | 62 | 61 | 248 | 4.56 |
|  | *hhhl* | 0 | 187 | 0 | 61 | 248 | 5.17 |
|  | *hlll* | 0 | 61 | 0 | 187 | 248 | 4.08 |
|  | *mmmm* | 0 | 0 | 247 | 1 | 248 | 5.39 |
|  | DNA<br>concentration<br>evenness | High (*H*) | high (*h*) | medium (*m*) | low (*l*) | Total number of<br>OTUs | Shannon index |
|  |  | 5-60 ng/μl | 0.1-3.0 ng/μl | 0.009-0.09 ng/μl | 0.0001-0.008 ng/μl |  |  |
| Legs | *Hhml* | 69 | 63 | 63 | 53 | 248 | 4.21 |
|  | *hhhl* | 0 | 195 | 0 | 53 | 248 | 5.04 |
|  | *hlll* | 0 | 71 | 0 | 177 | 248 | 4.13 |
|  | *mmmm* | 0 | 0 | 238 | 10 | 248 | 5.32 |

#### Creation of mock-soups

– We used 1.0 μl aliquots of the appropriately diluted leg and body DNA extracts of the 248 input species to create eight mock soups, achieving different profiles of COI-marker-concentration evenness: *Hhml, hhhl, hlll*, and *mmmm*, where *H, h, m*, and *l* represent four different concentration levels (Fig. 1, Table 2). For instance, in the *Hhml* soups, approximately one-fourth of the input species were added at each concentration level (*H, h, m, l*), whereas in the *hlll* soup, three-quarters of the species were diluted to the low concentration level before being added. These soups thus represent eight bulk samples with different absolute DNA concentrations (*leg* vs. *body*) and species evennesses (*Hhml, Hhml, hhhl, hhhl*).

**Fig 1.**
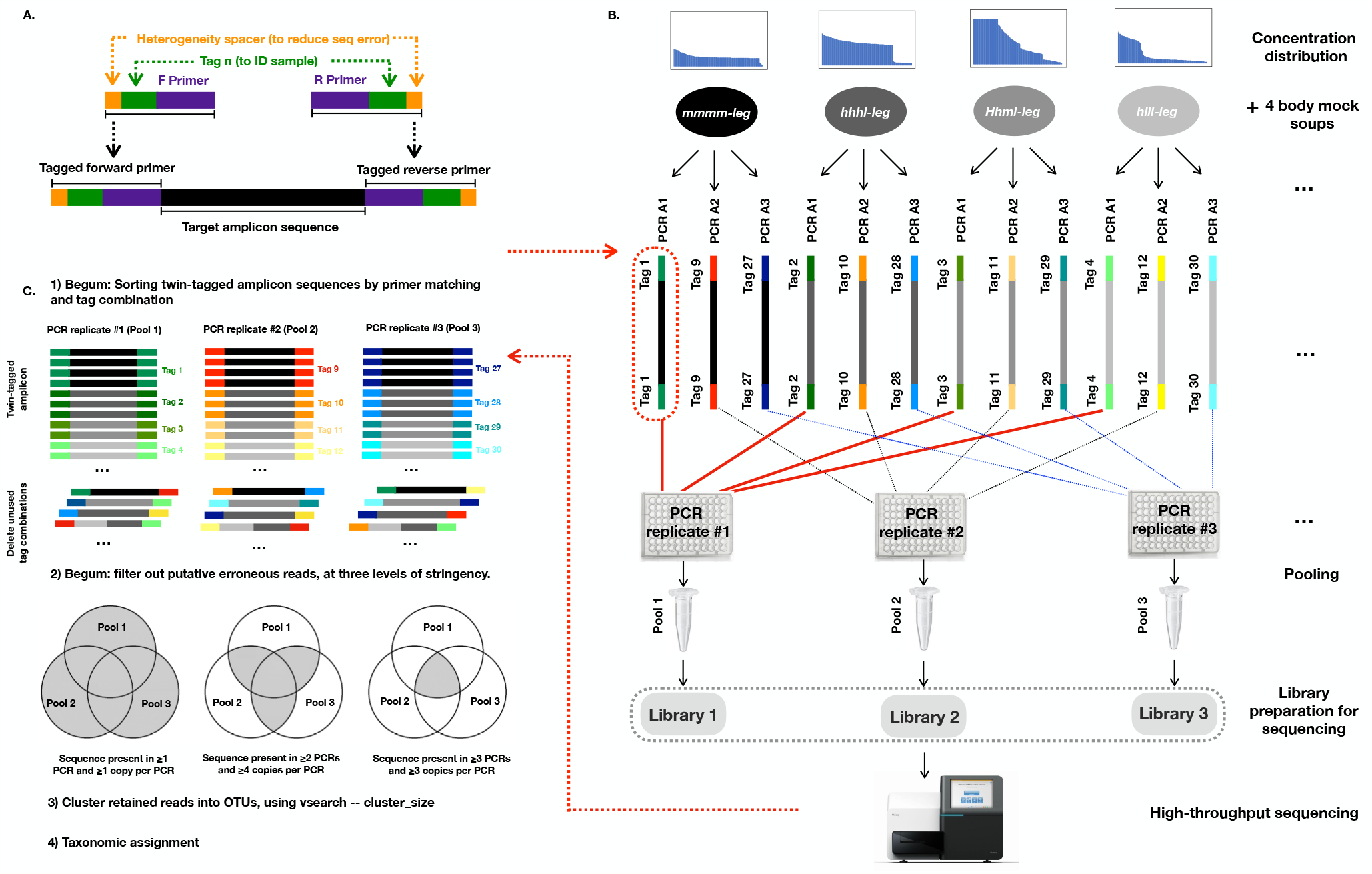
Schematic of study. **A**. Twin-tagged primers with heterogeneity spacers (above) and final amplicon structure (below). **B**. Each mock soup (e.g. *Hhml-leg*) was PCR-amplified three times (1, 2, 3) under a given PCR condition (A-H). Each of the three PCRs per soup used a different twin tag, following the *Begum* strategy. There were eight mock soups (*Hhml*/*hhhl*/*hlll/mmmm* X body/leg), where *H, h, m*, and *l* indicate different DNA concentrations (details in Figure 2). PCR replicates 1 from each of the eight mock soups were pooled into the first amplicon pool (solid red lines), PCR replicates 2 were pooled into the second amplicon pool (black dashes), and PCR replicates 3 were pooled into the third amplicon pool (blue dashes). The entire setup in B was repeated eight times for the eight PCR experiments (A-H), which thus generated (3 × 8 =) 24 sequencing libraries. **C**. Key steps of the *Begum* bioinformatic pipeline. For clarity, primers and heterogeneity spacers not shown. The complete PCR setup schematic, including positive and negative controls, is in S09.

### Primer tag design

For DNA metabarcoding, we also used the Leray-FolDegenRev primer set, which has been shown to result in a high recovery rate of arthropods from mixed DNA soups (Leray *et al*. 2013; Alberdi *et al*. 2018), and we used OligoTag (Coissac 2012) (Table S10) to design 100 unique tags of 7 nucleotides in length in which no nucleotide is repeated more than twice, all tag pairs differ by at least 3 nucleotides, no more than 3 G and C nucleotides are present, and none ends in either G or TT (to avoid homopolymers of GGG or TTT when concatenated to the Leray-FolDegenRev primers). We added one or two ‘heterogeneity spacer’ nucleotides to the 5’ end of the forward and reverse primers (De Barba *et al*. 2014; Fadrosh *et al*. 2014), which cause sets of amplicons to be sequenced out of phase on the Illumina plate, reducing basecalling errors. The total amplicon length including spacers, tags, primers, and markers was expected to be ∼382 bp. The primer sequences are listed in Table S10.

### PCR optimization

We ran test PCRs using the Leray-FolDegenRev primers with an annealing temperature (T_a_) gradient of 40 to 64°C. Based on gel-band strengths, we chose an ‘optimal’ T_a_ of 45.5°C (clear and unique band on an electrophoresis gel) and a ‘high’ T_a_ value of 51.5 °C (faint band) to compare their effects on species recovery.

We followed Murray, Coghlan and Bunce (2015) (see also Bohmann *et al*. 2018) and first ran the eight mock soups through qPCR to establish the correct dilution per soup so as to minimise PCR inhibition, to assess extraction-negative controls, and to estimate the minimum cycle number needed to amplify the target fragment across samples. Based on the qPCR amplifications, we diluted 6 of the 8 soups by 5, 10, or 50-fold to minimize inhibition (S06_Extended Methods), and we observed that the end of the exponential phase for all eight soups was achieved at or near 25 cycles, which we define here as the ‘optimal’ cycle number. To test the effect of PCR cycle number on species recovery, we also tested a ‘low’ cycle number of 21 (i.e. stopping amplification during the exponential phase), and a ‘high’ cycle number of 30 (i.e. amplifying into the plateau phase).

### PCR amplifications of mock soups

We metabarcoded the mock soups under 5 different PCR conditions:

**A, B**. Optimal T_a_ (45.5°C) and optimal PCR cycle number (25). A and B are technical replicates.

**C, D**. High T_a_ (51.5°C) and optimal PCR cycle number (25). C and D are technical replicates.

**E**. Optimal T_a_ (45.5°C) and low PCR cycle number (21).

**F** Optimal T_a_ (45.5°C) and high PCR cycle number (30).

**G, H**. Touchdown PCR (Leray & Knowlton 2015). 16 initial cycles: denaturation for 10 s at 95°C, annealing for 30 s at 62°C (−1°C per cycle), and extension for 60 s at 72°C, followed by 20 cycles at an annealing temperature of 46°C. G and H are technical replicates.

Following the *Begum* strategy, for each of the PCR conditions, each mock soup was PCR-amplified three times, each time with a different tag sequence on a different plate (Fig. 1). The same tag sequence was attached to the forward and reverse primers of a given PCR, which we call ‘twin-tagging’ (e.g. F1-R1, F2-R2,…), to allow detection and removal of tag-jumped sequences, which produce non-twinned tags (e.g. F1-R2, F2-R3,…). This lets us avoid assigning sequences to the wrong samples (Schnell, Bohmann & Gilbert 2015). In each PCR plate, we also included one positive control (with four insect species), three extraction-negative controls, and a row of PCR negative controls. PCR and tag setups are in Table S09.

### Ilumina high-throughput sequencing

Sequencing libraries were created with the NEXTflex Rapid DNA-Seq Kit for Illumina (Bioo Scientific Corp., Austin, USA), following manufacturer instructions. In total, we generated 24 sequencing libraries (= 8 PCR conditions (A-H) × 3 PCR replicates/condition) (Fig. 1), of which 18 were sequenced in one run of Illumina’s V3 300 PE kit on a MiSeq at the Southwest Biodiversity Institute, Regional Instrument Center in Kunming. The 6 libraries from PCR conditions G and H were sequenced on a different run with the same kit type.

### Data processing

We removed adapter sequences, trimmed low-quality nucleotides, and merged read-pairs with default parameters in *fastp* 0.20.1 (Chen *et al*. 2018). To allow fair comparion across PCR conditions, we subsampled 350,000 reads from each of the 24 libraries to achieve the same depth.

*Begum* is available at https://github.com/shyamsg/Begum (accessed 13 Nov 2020). First, we used *Begum*’s *sort*.*py* (`-pm 2 -tm 1`) to demultiplex sequences by primers and tags, add the sample information to header lines, and strip the spacer, tag, and primer sequences. *Sort*.*py* reports the number of sequences that have novel tag combinations, representing tag-jumping events (mean 3.87%). We then used *Begum*’s *filter*.*py* to remove sequences < 300 bp and to filter out false-positive (erroneous) sequences (PCR and sequencing errors, chimeras, low-level contamination). We filtered at twelve levels of stringency: ≥1-3 PCRs × ≥1-4 copies per PCR. For instance, ≥1 PCR and ≥1 copy represents no filtering, as this allows even single sequences that appear in only one PCR (0_0_1, 0_1_0, 1_0_0), while ≥2 PCRs and ≥4 copies is moderately stringent, as it allows only sequences that appear in at least 2 PCRs with at least 4 copies each (e.g. 32_4_0 but not 32_2_0).

We used *vsearch* 2.15.0 (Rognes *et al*. 2016) to remove *de novo* chimeras (`--uchime_denovo`) and to produce a fasta file of representative sequences for 97% similarity Operational Taxonomic Units (OTUs, `--cluster_size`) and a sample × OTU table (`--otutabout`). We assigned high-level taxonomies to the OTUs using *vsearch* (`--sintax`) on the MIDORI COI database (Leray *et al*. 2018) and only retained the OTUs assigned to Arthropoda with probability ≥ 0.80. In *R* 4.0.0 (R Core Team, 2018), we set all cells in the OTU tables that contained only one read to 0 and removed the control samples.

### Metabarcoding efficiency

#### False-negative and false-positive frequencies

– For each of the eight mock-soups (Table 2), eight PCRs (A-H), and 12 Begum filtering stringencies (Tables 3, S05), we used *vsearch* (`--usearch_global`) to match the OTUs against the 248 input species and the four positive-control species (S07_MTBFAS.fasta), and we removed any OTUs in the mock soups that matched a positive-control species. False negatives (dropouts) are defined as any of the 248 input species that failed to be matched by one or more OTUs at ≥ 97% similarity, and false positives are defined as OTUs that matched to no input species at ≥97% similarity. For clarity, we only display results from the *mmmm_body* soups; all results can be accessed in the provided script.

**Table 3.** Species-recovery success by three *Begum* filtering stringency levels and five PCR conditions, using the *mmmm_body* soup. Recovered species are OTUs that match one of the 248 reference species at ≥97% similarity. False negatives (dropouts) are defined as reference species that fail to be matched by any OTU at ≥97% similarity. False-positive sequences are defined as OTUs that fail to match any reference species at ≥97% similarity. *Begum* filtering strongly reduces false-positive frequencies (dark-to light-orange cells) at the cost of a small rise in dropout frequency, especially for optimal PCR conditions (PCRs A, B, E) (light-to dark-blue cells). With non-optimal PCR conditions (PCRs C, D, F, G, H), the trade-off is stronger; filtering to reduce false positives strongly increases dropouts (blue cells are darker on the right hand side of the table). See *Effects of PCR condition and* Begum *filtering* for more details. Table **S05** shows the same information for all twelve *Begum* stringency levels.

| Begum filtering parameters | PCR conditions |  |  |  |  |  |  |  |
| --- | --- | --- | --- | --- | --- | --- | --- | --- |
|  | Optimum T<sub>a</sub><br>+ optimum cycle<br>number<br>(45.5°C, 25) |  | Optimum T<sub>a</sub><br>+ low cycle<br>number<br>(45.5°C, 21) | High T<sub>a</sub> + optimum<br>cycle number<br>(51.5°C, 25) |  | Optimum T<sub>a</sub><br>+ high cycle<br>number<br>(45.5°C, 30) | Touchdown PCR<br>(62-46°C,<br>16+20 cycles) |  |
| Present in $\geq 1$ PCR replicate with $\geq 1$ copies per PCR (no filtering) | A | B | E | C | D | F | G | H |
| Recovered species: OTUs matched to Refs ($\geq 97\%$ similarity) | 241 | 243 | 243 | 240 | 239 | 241 | 236 | 235 |
| False-negative sequences (dropouts) | 7 | 5 | 5 | 8 | 9 | 7 | 12 | 13 |
| **% False negatives (dropouts)** | **3%** | **2%** | **2%** | **3%** | **4%** | **3%** | **5%** | **5%** |
| False-positive sequences | 165 | 161 | 181 | 186 | 132 | 179 | 99 | 124 |
| **% False positives** | **67%** | **65%** | **73%** | **75%** | **53%** | **72%** | **40%** | **50%** |
| Present in $\geq 2$ PCR replicates with $\geq 4$ copies per PCR | A | B | E | C | D | F | G | H |
| Recovered species: OTUs matched to Refs ($\geq 97\%$ similarity) | 234 | 229 | 232 | 217 | 204 | 203 | 161 | 171 |
| False-negative sequences (dropouts) | 14 | 19 | 16 | 31 | 44 | 45 | 87 | 77 |
| **% False negatives (dropouts)** | **6%** | **8%** | **7%** | **13%** | **18%** | **18%** | **35%** | **31%** |
| False-positive sequences | 5 | 5 | 7 | 3 | 2 | 3 | 3 | 4 |
| **% False positives** | **2%** | **2%** | **3%** | **1%** | **1%** | **1%** | **1%** | **2%** |
| Present in $\geq 3$ PCR replicates with $\geq 3$ copies per PCR | A | B | E | C | D | F | G | H |
| Recovered species: OTUs matched to Refs ($\geq 97\%$ similarity) | 231 | 228 | 235 | 198 | 192 | 183 | 126 | 136 |
| False-negative sequences (dropouts) | 17 | 20 | 13 | 50 | 56 | 65 | 122 | 112 |
| **% False negatives (dropouts)** | **7%** | **8%** | **5%** | **20%** | **23%** | **26%** | **49%** | **45%** |
| False-positive sequences | 4 | 4 | 6 | 2 | 2 | 3 | 2 | 1 |
| **% False positives** | **2%** | **2%** | **2%** | **1%** | **1%** | **1%** | **1%** | **0%** |

#### Input DNA concentration and evenness and PCR conditions

– We used non-metric multidimensional scaling (NMDS) (`metaMDS(distance=“jaccard”, binary=FALSE)`) in {vegan} 2.5-6 (Oksanen *et al*. 2017) to visualise differences in OTU composition across the eight mock-soups per PCR condition (Fig. 1, Table 2). We evaluated the effects of species evenness on species recovery by using a linear mixed-effects model to regress the number of recovered input species on each mock soup’s Shannon diversity (Table 2), `lme4::lmer(OTUs∼Evenness+(1|PCR)` (Bates *et al*. 2015). Finally, we evaluated the information content of OTU size (number of reads) by linearly regressing input DNA concentration on OTU size.

#### Taxonomic bias

– To visualize the effects of PCR conditions on taxonomic amplification bias, we used {metacoder} 0.3.4 (Foster, Sharpton & Grunwald 2017) to pairwise-compare the compositions of the *mmmm_body* soup under different PCR conditions.

#### Tag-bias test

We took advantage of the paired technical replicates in PCRs A&B, C&D, and G&H (Table 3) to test for tag bias. For instance, we used the same eight tags in PCRs A1 and B1, and this pair should therefore return very similar communities. In contrast, non-matching pairs (e.g. A1 and B2, A2 and B1) used different tags and, if there is tag bias, should therefore return differing communities. For each PCR set (A&B, C&D, G&H), we generated NMDS ordinations and used `vegan::protest` to calculate the mean Procrustes correlation coefficients of PCR pairs that used the same tags (n = 3) and different tags (n = 12).

## Results

The 18 libraries containing PCR sets A-F yielded 7,139,290 total paired-end reads, mean 396,627, and the 6 libraries of PCR sets G&H yielded 6,356,655 paired-end reads, mean 1,059,442. Each sample (e.g. *Hhml_body* + *PCR_A*) was sequenced in three libraries (Figs. 1, S5) and thus was represented by a mean of 132,209 reads (=396,627 mean reads per library X 3 PCRs / 9 samples per library. Each library contains 8 mock soups + 1 positive control.) in PCR sets A-F and 353,147 reads in PCR sets G and H.

### Effects of PCR condition and Begum *filtering*

Optimal and near-optimal PCR conditions (PCRs A, B, E) achieved lower false-negative (dropout) frequencies than did non-optimal PCRs (high T_a_, high cycle number, or Touchdown) (PCRs C, D, F, G, H) (Table 3, S05).

With no *Begum* filtering (≥1 PCR & ≥1 copy), false-positive OTUs were abundant, approaching the number of true OTUs (101-187 false-positive OTUs versus 248 true OTUs) (Table 3, S05). Applying *Begum* filtering at different stringency levels reduced the number of false-positive sequences by 3 to 90 times, and even eliminated them in one case. The cost of filtering was a greater loss of true OTUs but only by a small absolute amount in the optimal PCRs (A, B, E), rising from a dropout frequency of ∼2% in the nonfiltered case to ∼4-6% under all but the two most stringent filtering levels, where dropout frequencies were 5-11% (≥3 PCRs & ≥3 or 4 copies/PCR). In contrast, in the non-optimal PCRs (C, D, F, G, H), *Begum* filtering caused dropout frequencies to rise to much higher levels (5-55%). In short, it is possible to combine wet-lab and bioinformatic protocols to simultaneously reduce false-positive and false-negative errors.

### Effects of input-DNA absolute and relative concentrations on OTU recovery

Altering the relative (*Hhml, hhhl, hlll*, and *mmmm*) and absolute (body, leg) input DNA concentrations created quantitative compositional differences in the OTU tables, as shown by NMDS ordination (Fig. 2). Soup *hlll*, with the most uneven distribution of input DNA concentrations (Table 2), recovered the fewest OTUs (Fig. 2). The same effect was seen by regressing the number of recovered OTUs on species evenness (Fig. S01).

**Fig. 2.**
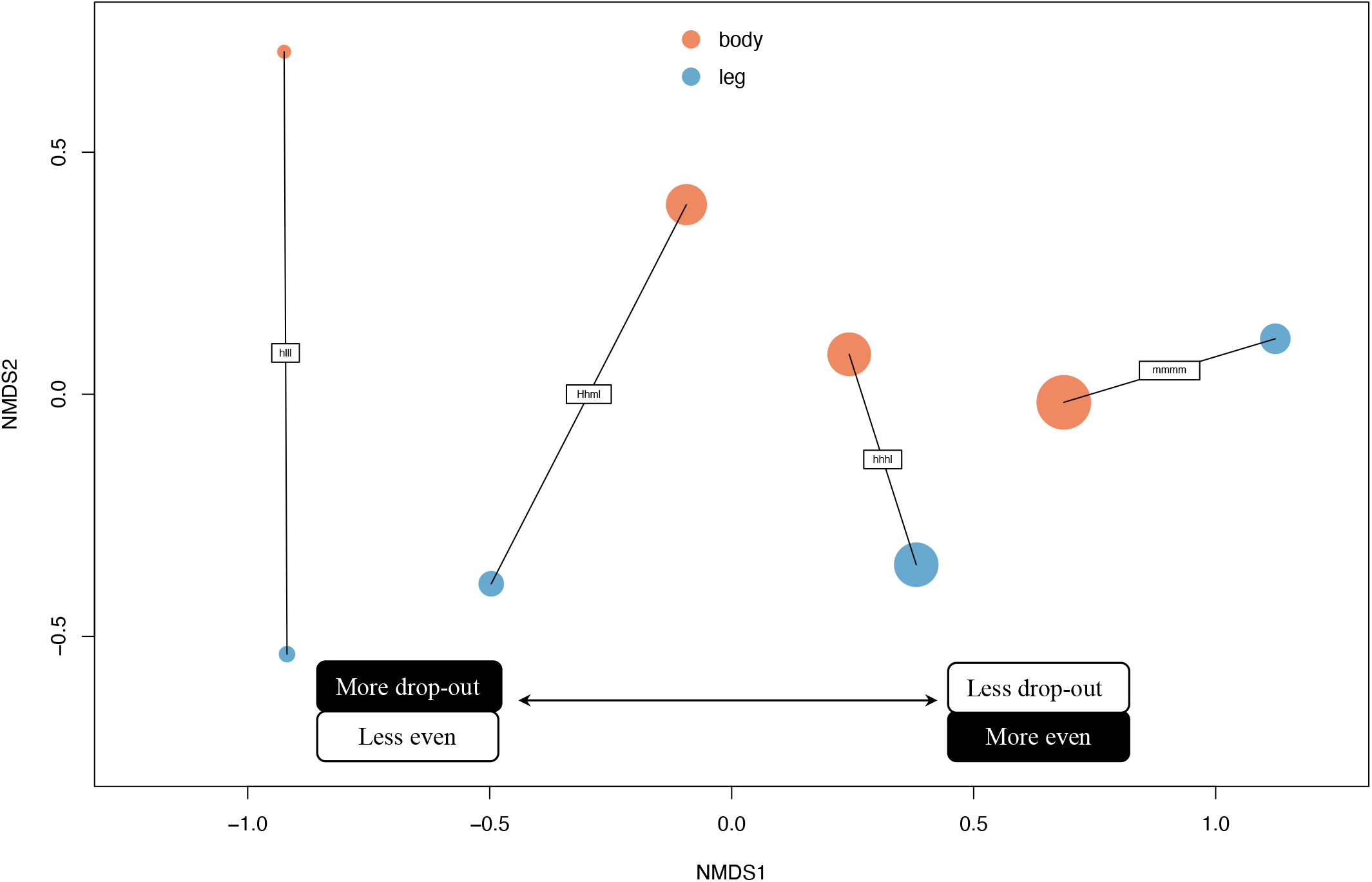
Non-metric multidimensional scaling (NMDS) ordination of eight mock soups, which differ in absolute (Body/Leg) and relative (*Hhml, hhhl, hlll*, and *mmmm*) DNA concentrations of the input species (Table 2). Shown here is the output from the PCR A condition: optimum annealing temperature T_a_ (45.5 C) and cycle number (25), at *Begum* filtering stringency ≥2 PCRs, ≥4 copies/PCR (Table 3). Point size is scaled to the number of recovered OTUs. Species recovery is lower in samples with more uneven species frequencies (e.g. *hlll*) and, to a lesser extent, lower absolute DNA input (leg).

As expected, OTU size does a poor job of recovering information on input DNA amount per species (Fig. S02). Although there are positive relationships between OTU size and DNA concentrations, the slope of the relationship differs depending on species frequencies (*Hhml* vs. *hhhl* vs. *hlll*) and source tissues (leg vs. body), which reflects the action of multiple species-specific biases along the metabarcoding pipeline (McLaren, Willis & Callahan 2019). This interaction effect precludes the fitting of a robust model that relates OTU size to DNA concentration, since species-frequency and source-tissue information are not known *a priori*.

### Taxonomic amplification bias

Optimal PCR conditions (PCRs A, B, E) produce larger OTUs than do non-optimal PCR conditions (PCRs C, D, F, G, H), especially for Hymenoptera, Araneae, and Hemiptera (Fig. 4). These are the taxa that are at higher risk of failing to be detected by the Leray-FolDegenRev primers under sub-optimal PCR conditions.

**Fig. 3.**
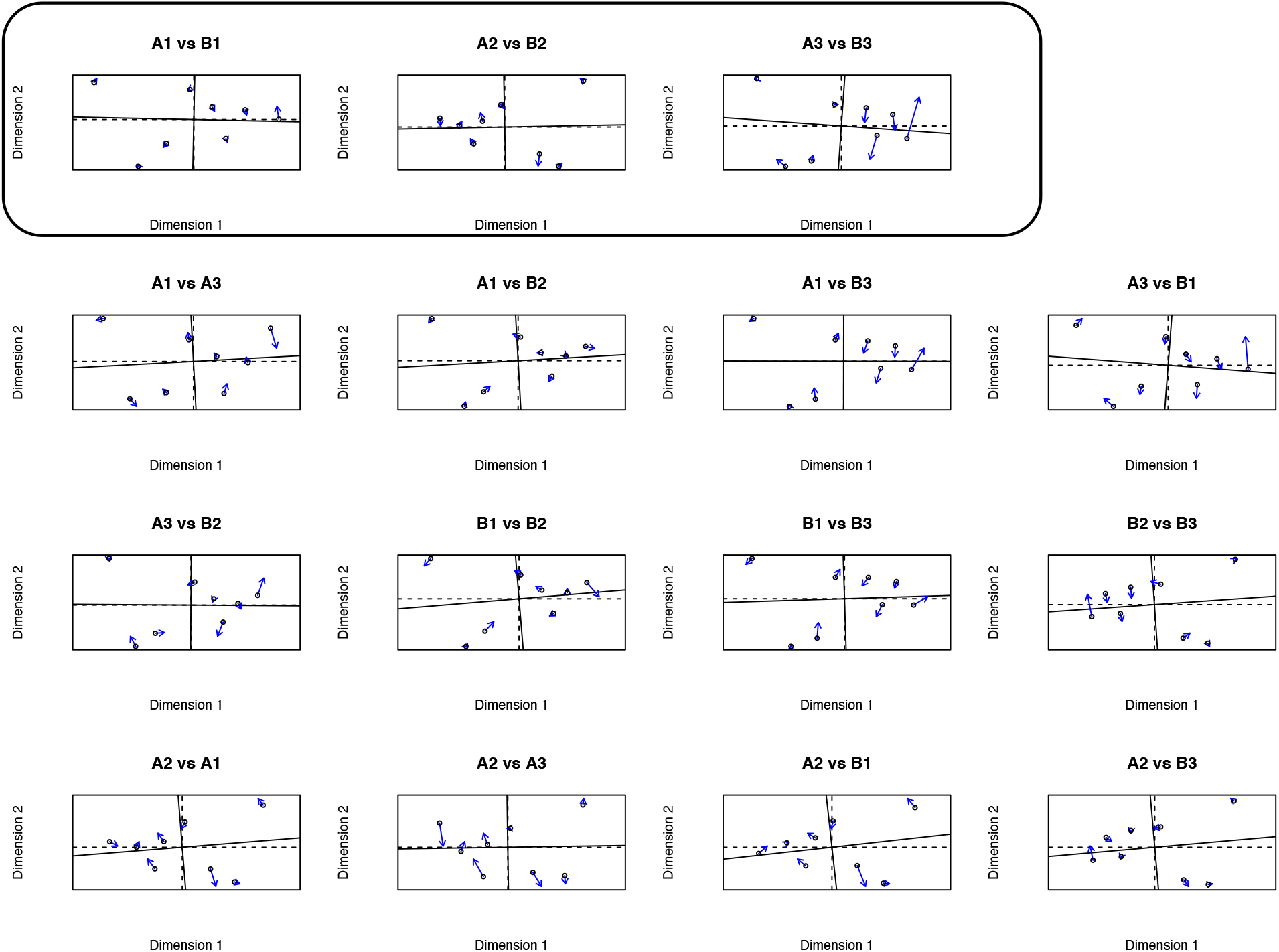
Test for tag bias in the mock soups amplified at optimum annealing temperature T_a_ (45.5 °C) and optimum cycle number (25) (PCRs A and B). All pairwise Procrustes correlations of PCRs A and B. The top row (box) displays the three same-tag pairwise correlations. The other rows display the 12 different-tag pairwise correlations. If there is tag bias during PCR, the top row should show a greater degree of similarity. However, mean correlations are not significantly different between same-tag and different-tag ordinations (Mean of same-tag correlations: 0.99 ± 0.007 SD, n = 3. Mean of different-tag correlations: 0.98 ± 0.009 SD, n = 12. p=0.046, df=3.9, Welch’s t-test). In Supplementary Information, we show the results for the high T_a_ (PCRs C & D) and Touchdown treatments (PCRs G & H).

**Fig. 4.**
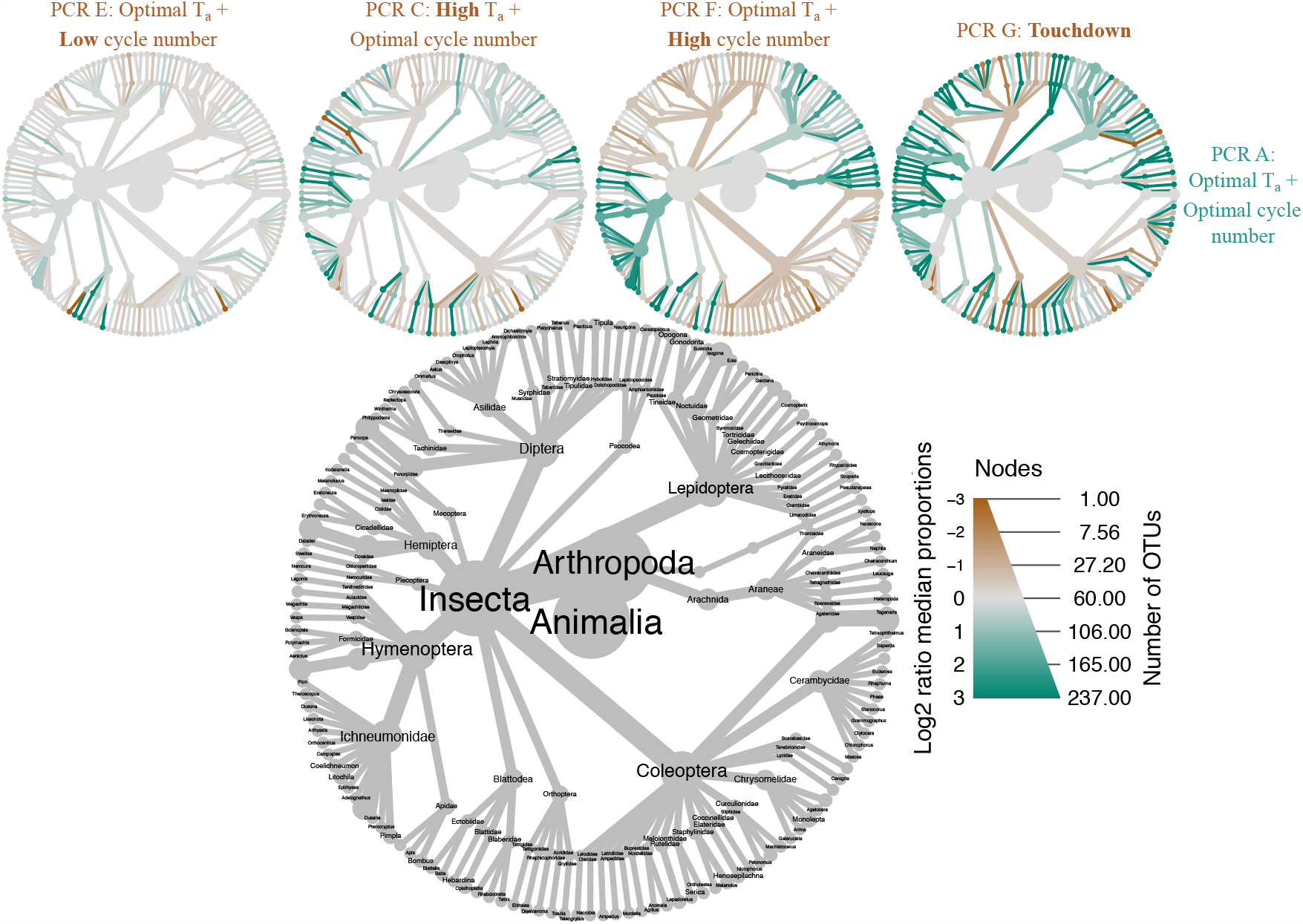
Taxonomic amplification bias of non-optimal PCR conditions. Pairwise-comparison heat trees of PCRs E, C, F, & G versus PCR A (Table 3). Green branches indicate that PCR A (right side) produced relatively larger OTUs in those taxa. Brown branches indicate that PCR A produced smaller OTUs. Grey branches indicate similar OTU sizes. There are, on balance, more dark green branches than dark brown branches in the three heat trees that compare PCRs C, F, and G (sub-optimal) with PCR A (optimal), and the green branches are concentrated in the Araneae, Hymenoptera, and Lepidoptera, suggesting that these are the taxa at higher risk of failing to be detected by Leray-FolDegenRev primers under sub-optimal PCR conditions. Shown here are the *mmmmbody* soups, at *Begum* filtering stringency ≥2 PCRs, ≥4 copies per PCR.

### Tag-bias test

We found no evidence for tag bias in PCR amplification. For instance, under optimal PCR conditions (A & B), pairs using the same tags (A1/B1, A2/B2, A3/B3) and pairs using different tags (e.g. A1/B2, A2/B1, A3/B2) both generated almost identical NMDS ordinations (Fig. 3). Under non-optimal PCRs, we still found no evidence for tag bias, even though at high annealing temperatures, it is theoretically possible that some tag sequences might be more likely to aid primer annealing (Fig. S03, S04). Note that we did not correct the p-values for three tests, underlining the lack of evidence for tag bias.

## Discussion

In this study, we tested our pipeline with eight mock soups that differed in their absolute and relative DNA concentrations of 248 arthropod taxa (Table 2, Fig. 2). We metabarcoded the soups under five different PCR conditions that varied annealing temperatures (T_a_) and PCR cycles (Table 3), and we used *Begum* to filter the OTUs under different stringencies (Fig. 1, Table 3). We define high efficiency in metabarcoding as recovering most of sample’s compositional and quantitative information, which in turn means that both false-negative and false-positive frequencies are low, that OTU size predicts biomass or abundance per species, and that any dropouts are spread evenly over the taxonomic range of the target taxon (here, Arthropoda). This pipeline can of course be applied to other taxa, with appropriate adjustments to primer design, length limits, taxonomic reference database, and positive controls.

Our results show that metabarcoding efficiency can be made high for the recovery of species presence-absence, but efficiency is low for the recovery of quantitative information. Efficiency increases when the annealing temperature and PCR cycle number are at the low ends of ranges currently reported in the literature for this primer pair (Table 3, Fig. 4). We recovered Elbrecht et al.’s (2017) finding that efficiency is higher when species evenness is higher (Fig. 2, S01), and we found that OTU sizes are a poor predictor of input genomic DNA, which confirms the conventional wisdom that OTU size is a poor predictor of species relative abundances (Fig. S02) (McLaren, Willis & Callahan 2019). Finally, we found no evidence for tag bias during PCR (Figs. 3, S03, S04).

### Co-designed wet-lab and bioinformatic methods to remove errors

The *Begum* workflow co-designs the wet-lab and bioinformatic components (Fig. 1) (Zepeda-Mendoza *et al*. 2016) to minimise multiple sources of error (Table 1). Aside from the use of qPCR to optimise PCR conditions, the wet-lab and bioinformatic components are designed to work together. Twin-tagging allows removal of tag jumps, which result in sample misassignments. Multiple, independent PCRs per sample allow removal of false-positive sequences caused by PCR and sequencing error and by low-level contamination, at the cost of only a small absolute rise in false-negative error (Tables 3, S05). qPCR optimization reduces false negatives caused by PCR runaway, PCR inhibition, and annealing failure (Tables 3, S05; Fig. 4). Moderate dilution appears to be a better solution for inhibition than is increasing cycle number, since the latter increases dropouts (Tables 3, S05). qPCR also allows extraction blanks to be screened for contamination. While size sorting (Elbrecht, Peinert & Leese 2017) should reduce false negatives caused by PCR runaway, the lower recovery of input species in the leg-only soups (Fig. 2) argues that large insects should be reduced to at least their heads, not their legs, for DNA extraction.

### Begum filtering and complex positive controls

Increasing the stringency of *Begum* filtering reduces false-positive sequences at the cost of increasing false-negatives (dropouts), although fortunately, this trade-off is weakened under optimal PCR conditions (Tables 3. S05). The choice of a filtering stringency level for a given study should be informed by complex positive-control samples and should take into account the study’s aims. If the aim is to detect a particular taxon, like an invasive pest, it is better to set stringency low to minimise dropout, whereas if the aim is to generate data for an occupancy model, it is better to set stringency high to minimise false positives. Positive controls should be made of diverse taxa not from the study area (Creedy, Ng & Vogler 2019) and span a range of concentrations. Alternatively, a suite of synthetic DNA sequences with appropriate primer binding regions could be used.

In metabarcoding pipelines, it is common to apply heuristic filters to remove false-positive sequences. For instance, small OTUs are commonly removed (http://evomics.org/wp-content/uploads/2016/01/phyloseq-Lab-01-Answers.html, accessed 11 Nov 2020). We did not do this because we wanted to isolate the effect of *Begum* filtering (and in fact we found that doing so slightly reduced species recovery). We did set to zero all cells in our OTU tables that contained only one read, and the only effect was to greatly reduce the number of false-positive sequences in the case when *Begum* filtering was not applied. Once any level of *Begum* filtering had been applied, those 1-read cells also disappeared (D. Yu, data not shown). Another common correction is to use the *R* package {lulu} (Frøslev *et al*. 2017) to combine ‘parent’ and ‘child’ OTUs that had failed to cluster. In this study, we could not do this because all input species had been included in all eight soups, which means that OTU co-occurrence could not be used to identify parent-child pairings.

### Future work

*Begum* uses occurrence in multiple, independent PCRs to identify and remove erroneous sequences. This contrasts with solutions such as DADA2 (Callahan *et al*. 2016) and UNOISE2 (Edgar 2016) that use only sequence quality data to remove erroneous sequences. Unique molecular identifiers (UMIs) are also a promising method for the removal of erroneous sequences (Fields *et al*. 2019). It should be possible to combine some of these methods in the future.

A second area of research is to improve the recovery of quantitative information. Spike-ins and UMIs can be part of the solution (Smets 2016; Hoshino & Inagaki 2017; Deagle *et al*. 2018; Tkacz, Hortala & Poole 2018; Ji *et al*. 2020), but they can only correct for sample-to-sample stochasticity and differences in total DNA mass across samples. Such corrections allow tracking of *within-species change across samples*, which means tracking how each individual species changes in abundance along a time series or environmental gradient. However, spike-ins and UMIs cannot be used to estimate *species relative abundances within a sample*, because spike-ins do not remove species biases in DNA-extraction and primer-binding efficiencies. Thus, we caution against the uncritical use of metabarcoding to identify major and minor diet components (e.g. Deagle *et al*. 2019). Fortunately, methods for estimating species relative abundances are being developed (Lang *et al*. 2019; Peel *et al*. 2019; Williamson, Hughes & Willis 2019).

## Acknowledgements

We thank Mr. Zongxu Li in South China Barcoding Center for help with arthropod selection and morphological identification. C.Y.Y., D.W.Y., W.X.Y., W.C. were supported by the Strategic Priority Research Program of the Chinese Academy of Sciences (XDA20050202),the National Natural Science Foundation of China (41661144002, 31670536, 31400470, 31500305), the Key Research Program of Frontier Sciences, CAS (QYZDY-SSW-SMC024), the Bureau of International Cooperation (GJHZ1754), the Ministry of Science and Technology of China (2012FY110800), the State Key Laboratory of Genetic Resources and Evolution (GREKF18-04) at the Kunming Insitute of Zoology, the University of East Anglia, and the University of Chinese Academy of Sciences. D.W.Y. was supported by a Leverhulme Trust Research Fellowship. K.B. was supported by the Danish Council for Independent Research (DFF-5051-00140).

## Author contributions

D.Y. and C.Y.Y. designed the project; C.Y.Y and K.B. designed the laboratory protocol; C.Y.Y and W.C conducted the laboratory work; Z.L.D. performed the library building and Miseq sequencing; N.W. prepared the primer and tag design Excel spreadsheet; S.G. wrote *Begum*; X.Y.W wrote additional programs; D.Y. and C.Y.Y. wrote the bioinformatics pipeline and performed data analysis; D.Y. wrote the first draft of the paper, and C.Y.Y. and K.B. contributed revisions.

## Data Availability

Supplementary Information includes a tutorial with a reduced sequence dataset and simplified scripts (PCR B only, 253 MB). We have archived the full dataset (∼10 GB), with reference files, folder structure, output files, and scripts. To run the scripts, remove the included output files listed in the DataDryad readme. The archive is available for reviewer download at: https://datadryad.org/stash/share/_XecP4iDVnM2uFmFyCcEpV3bD58swl8lIuzc-2ZDMNc The scripts are kept updated at https://github.com/dougwyu/BiodiversitySoupII.

---

**Fig. S01.**
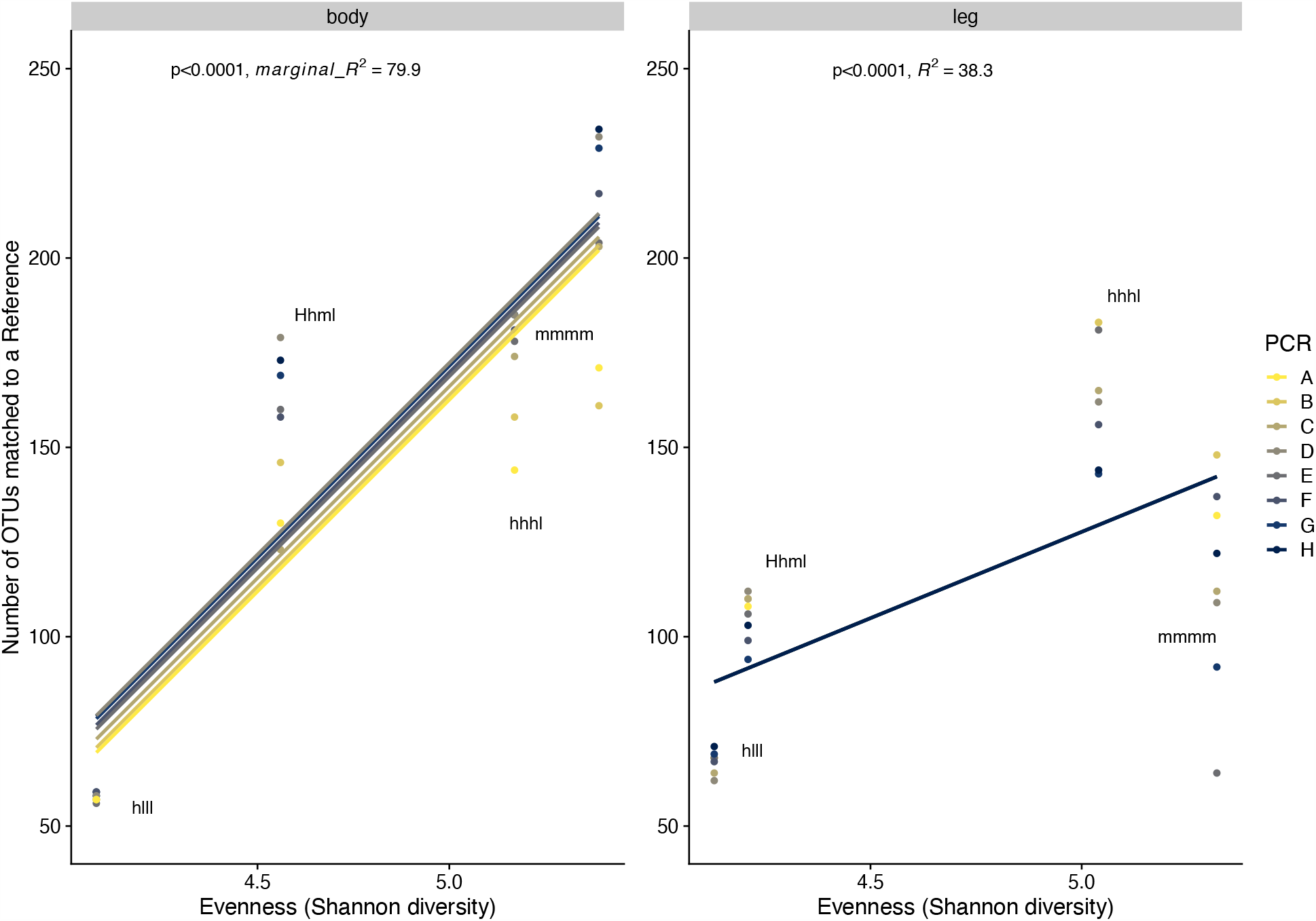
Effect of species evenness on OTU recovery. More input species are recovered with greater species evenness, as measured by Shannon diversity in Table 2. Shown here are the results from *Begum* filtering stringency of ≥2 PCR replicates & ≥4 reads. Body: summary statistics from mixed-effects model `lme4::lmer(OTUs∼Shannon+(1|PCR)`. Marginal R^2^ is the fixed-effect contribution `(MuMin::r.squaredGLMM)`. Leg: summary statistics from linear model.

**Fig. S02.**
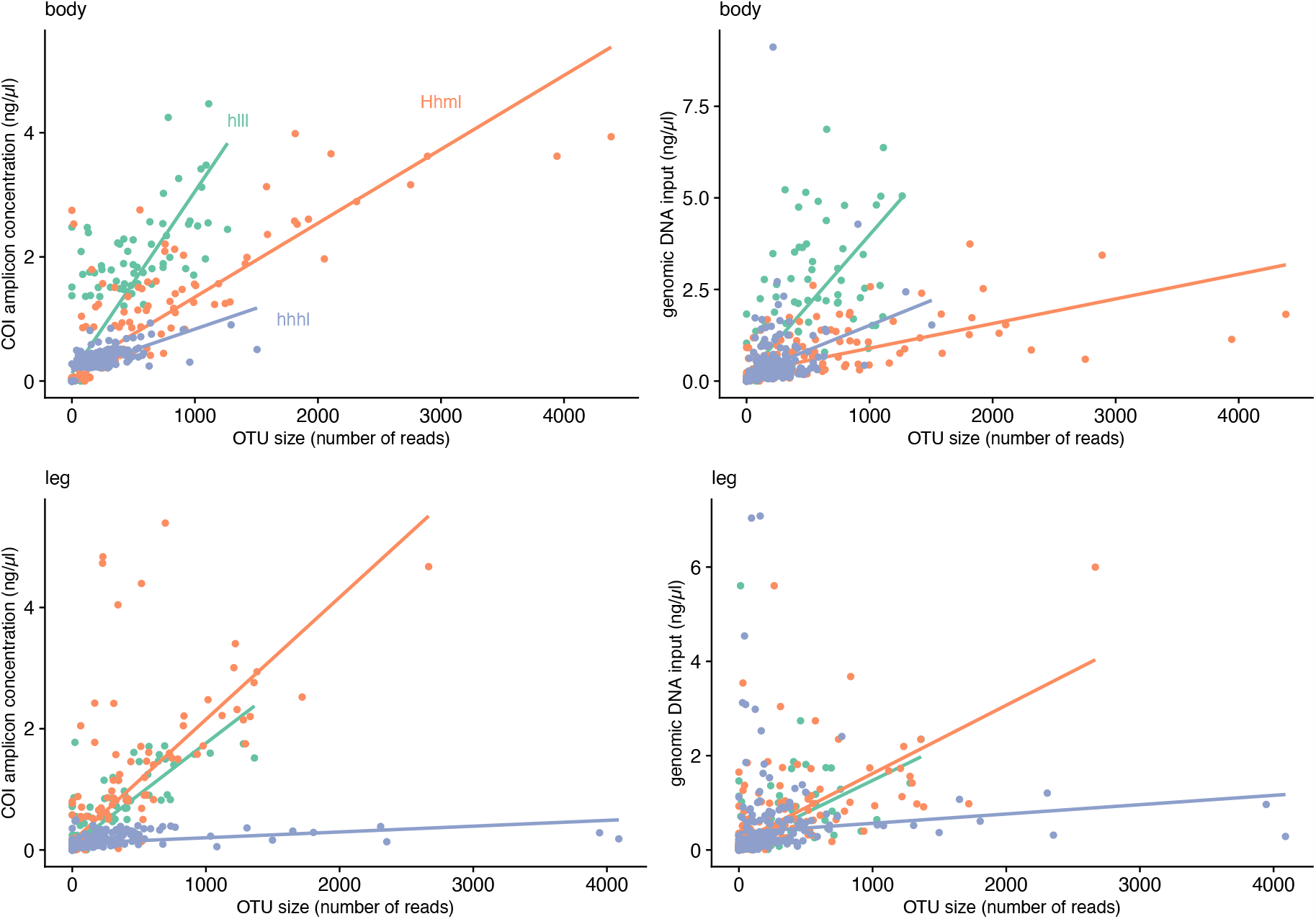
Recovery of species relative abundance information from read number. Each data point is one input species; fitted lines are linear regressions of input DNA concentration (*left*: COI; *right*: genomic DNA; *top*: body; *bottom*: leg) on OTU size (number of reads). The linear relationships between input DNA concentration and OTU size vary significantly by soup (*hlll, Hhml, hhhl*) (`lm(input_DNA_conc ∼ OTU_size + soup + OTU_size:soup`), all interaction-term *p*<0.0001). Shown is the dataset from PCR A after *Begum* filtering stringency of ≥2 PCR replicates & ≥4 reads (Table 3). The *mmmm* soups were omitted because they have almost no variance.

**Fig. S03.**
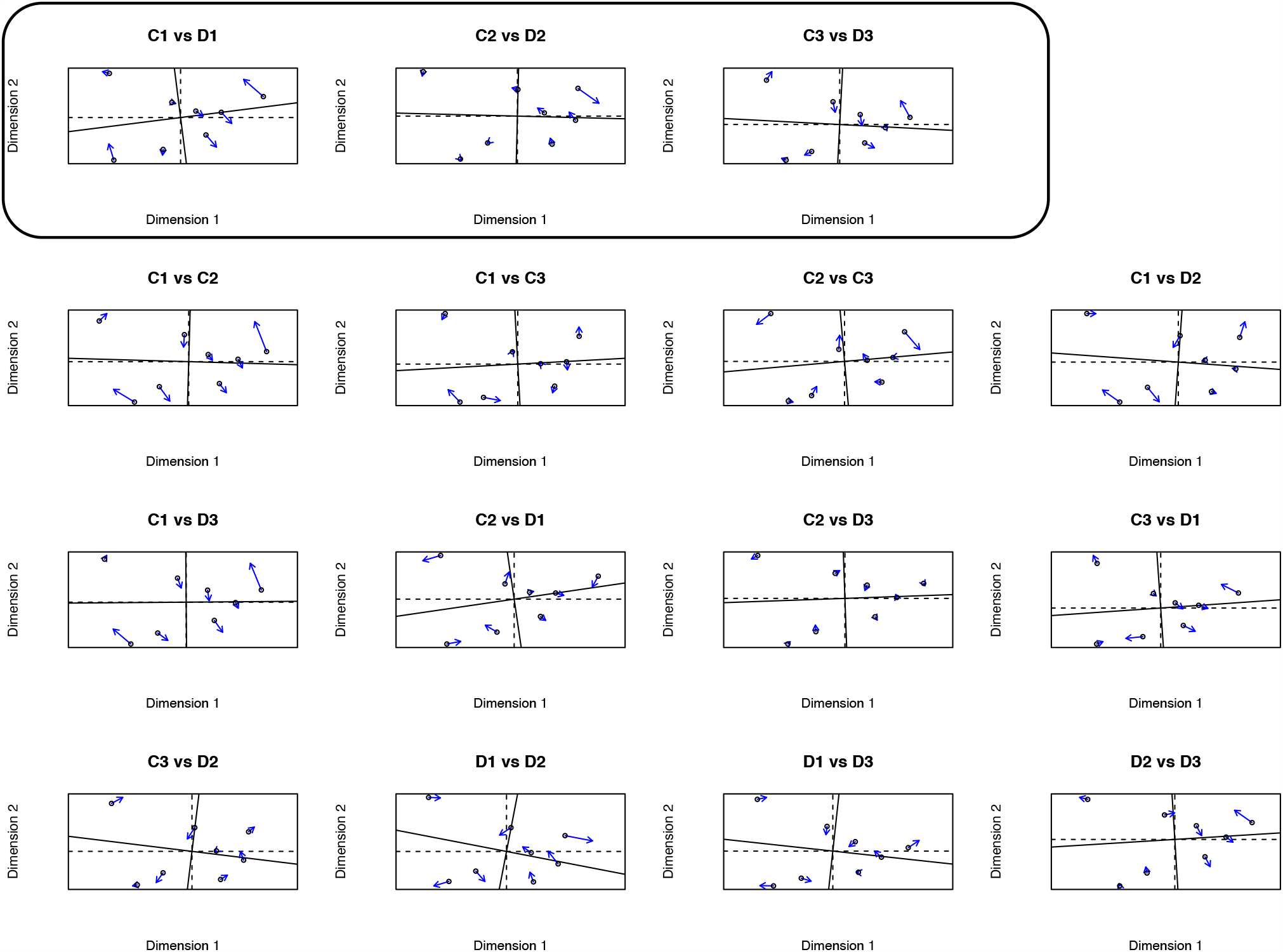
Test for tag bias in the mock soups amplified at high annealing temperature T_a_ (51.5 °C) and optimum cycle number (25) (PCRs C and D). All pairwise Procrustes comparisons of PCRs C and D. The top row (box) displays the three same-tag pairwise correlations. The other rows display the 12 different-tag pairwise correlations. If there is tag bias during PCR, the top row should show a greater degree of similarity. However, mean correlations are not significantly different between same-tag and different-tag ordinations (Mean of same-tag correlations: 0.98 ± 0.019 SD, n = 3; mean of different-tag correlations: 0.98 ± 0.011 SD, n = 12. P=0.352, df=17.8, Welch’s t-test).

**Fig. S04.**
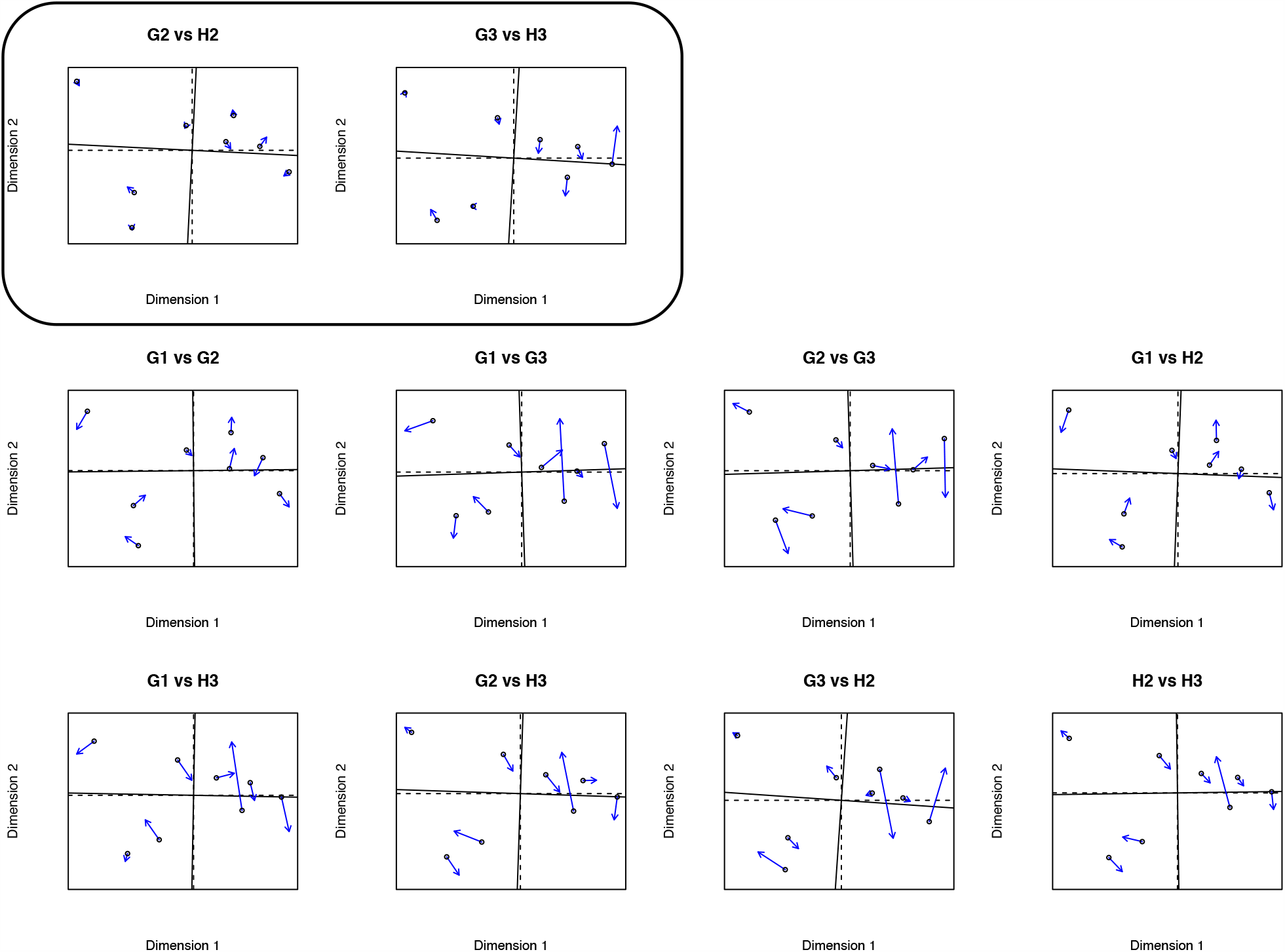
Test for tag bias in the mock soups amplified with the Touchdown PCR protocol (G and H). All pairwise Procrustes comparisons of PCRs G and H. The top row (box) displays the two pairwise comparisons between the technical replicates. The other rows display all the different-tag pairwise correlations. If there is tag bias during PCR, the top row should show a greater degree of similarity. Mean correlations are not significantly different between same-tag and different-tag ordinations, but we note that only two same-tag comparisons could be made (Mean of same-tag correlations: 0.98 ± 0.019 SD, n = 2; mean of different-tag correlations: 0.91 ± 0.033 SD, n = 8. P=0.072, df=2.9, Welch’s t-test).

**Table S05.**
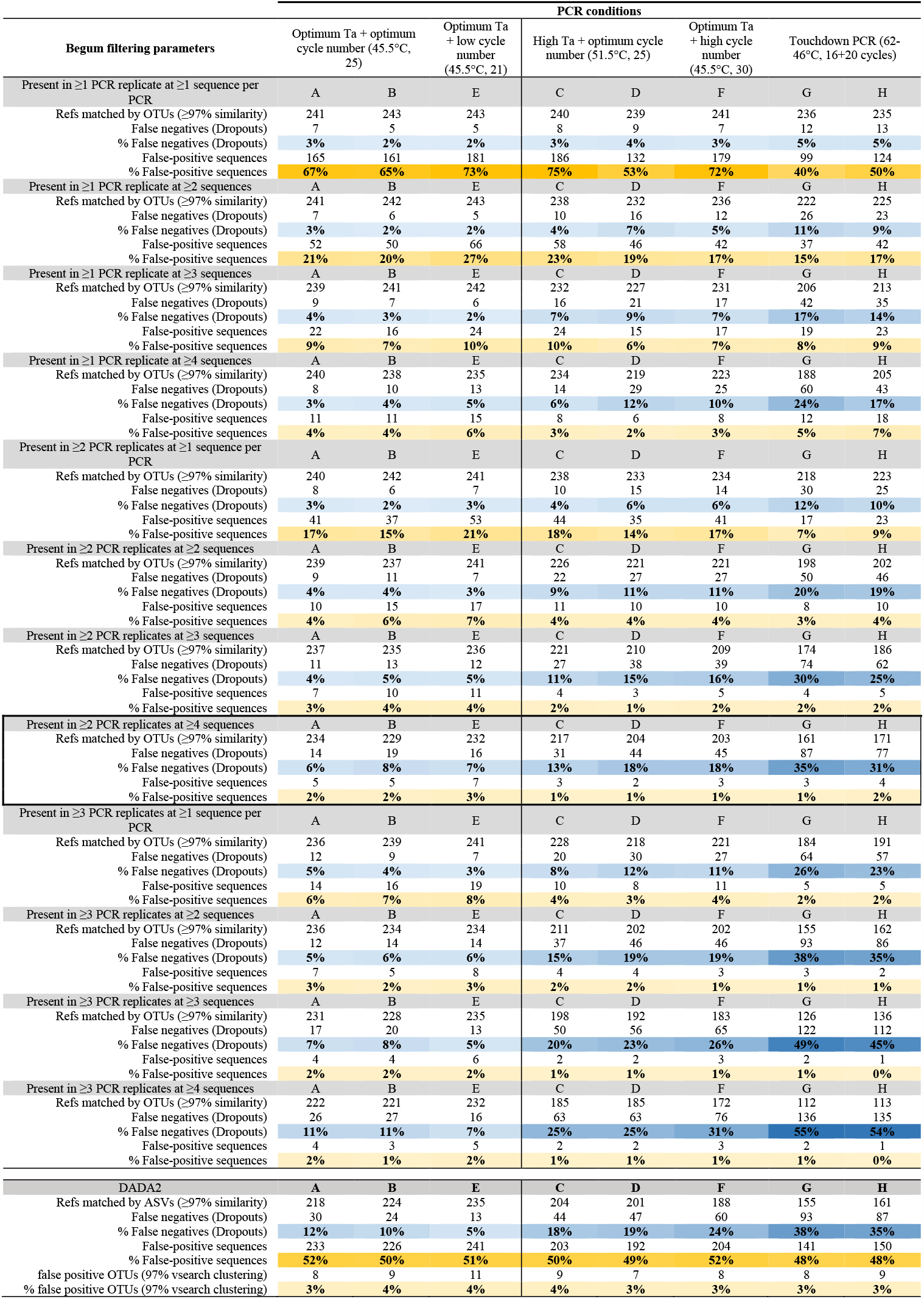
Species-recovery success by 12 *Begum* filtering stringency levels and five PCR conditions, compared with DADA2, using the *mmmm_body* soup. Recovered species are OTUs that match one of the 248 reference species at ≥97% similarity. False negatives (dropouts) are defined as reference species that fail to be matched by any OTU at ≥97% similarity. False-positive sequences are defined as OTUs that fail to match any reference species at ≥97% similarity. *Begum* filtering strongly reduces false-positive frequencies (transition from dark-to light-orange cells) at the cost of a smaller rise in dropout frequency (light-to dark-blue cells), especially for optimal PCR conditions (PCRs A, B, E). With non-optimal PCR conditions (PCRs C, D, F, G, H), the trade-off is stronger; filtering to reduce false positives strongly increases dropouts (blue cells are darker on the right hand side of the table). See *Effects of PCR condition and* Begum *filtering* for more details. An Excel worksheet version of this table is in Supplementary Information. Most analyses in *Results* were carried out at *Begum* stringency level ‘Present in ≥2 PCR replicates at ≥4 sequences’ (box).

